# Phylogenomic and phenotypic analyses highlight the diversity of antibiotic resistance and virulence in both human and non-human *Acinetobacter baumannii*

**DOI:** 10.1101/2023.12.13.571483

**Authors:** Ellen M. E. Sykes, Valeria Mateo-Estrada, Raelene Engelberg, Anna Muzaleva, George Zhanel, Jeremy Dettman, Julie Chapados, Suzanne Gerdis, Ömer Akineden, Izhar U.H. Khan, Santiago Castillo-Ramírez, Ayush Kumar

**Author notes:** Address correspondence to Ayush Kumar;, Ellen M. E. Sykes;, Valeria Mateo-Estrada;, Raelene, Engelberg;, Anna Muzaleva;, George Zhanel;, Jeremy Dettman;, Julie Chapados;, Suzanne Gerdis;, Ömer Akineden;, Izhar U. H. Khan;, Santiago Castillo-Ramírez;, Ayush Kumar.

## Abstract

*Acinetobacter baumannii* is a Gram-negative opportunistic pathogen causing infections of the immunocompromised. With a high incidence of muti-drug resistance, carbapenem-resistant *A. baumannii* is as a priority 1 pathogen designated by the WHO. The current literature has expertly characterized clinical isolates of *A. baumannii.* As the challenge of these infections has recently been classified as a One Health issue, we set out to explore the diversity of isolates from human and non-clinical sources such as agricultural surface water, urban streams, various effluents from wastewater-treatment plants and tank milk; and, importantly, these isolates came from a wide geographic distribution. Phylogenomic analysis considering almost 200 isolates showed that our diverse set is well-differentiated from the main international clones of *A. baumannii*. We discovered novel sequence types in both hospital and non-clinical settings, and five strains that overexpress the RND efflux pump *adeIJK* without changes in susceptibility. Further, we detected a *bla*_ADC-79_ in a non-human isolate despite its sensitivity to all antibiotics. There was no significant differentiation between the virulence profiles of clinical and non-clinical isolates in the *Galleria mellonella* insect model of virulence suggesting that virulence is neither dependent on geographic origin nor isolation source. Detection of antibiotic resistance and virulence genes in non-human strains suggests that these isolates may act as a genetic reservoir for clinical strains. This endorses the notion that in order to combat multi-drug resistant infection caused by *A. baumannii,* a One Health approach is required, and a deeper understanding of non-clinical strains must be achieved.

**Importance:** The global crisis of antibiotic resistance is a silent one. More and more bacteria are becoming resistant to all antibiotics available for treatment, leaving no options remaining. This includes *Acinetobacter baumannii.* This Gram-negative opportunistic pathogen shows a high frequency of multi-drug resistance, and many strains are resistant to last-resort drugs carbapenem and colistin. Research has focused on strains of clinical origin, but there is a knowledge gap regarding virulence traits, particularly, how *A. baumannii* become the notorious pathogen of today. Antibiotic resistance and virulence genes have been detected in strains from animals, and environmental locations such as grass and soil. As such, *A. baumannii* is a One Health concern which includes the health of humans, animals and the environment. Thus, in order to truly combat the antibiotic resistance crisis, we need to understand antibiotic resistance and virulence gene reservoirs of this pathogen under the One Health continuum.

**Repositories:** NCBI GenBank Accession numbers: Bioproject PRJNA819071, Biosamples SAMN26898552 - SAMN26898587.

## Introduction

*Acinetobacter baumannii* is as a Gram-negative hospital-acquired opportunistic pathogen. It causes ventilator-associated pneumonia, urinary tract and wound infections and can cause bacteremia in cases of treatment failure (1). Community-associated infections have been reported as rare but serious (2, 3). Additionally, carbapenem-resistant *A. baumannii* has been labelled a priority 1 pathogen by the World Health Organization (4) based on the inability to cure such infections. Due to the COVID-19 pandemic, the number of *A. baumannii* infections increased (5) and there is a high frequency of multi-drug resistance (MDR); upwards of 70% of strains (1). Presence of Antibiotic Resistance Genes (ARGs) and virulence genes (VGs) are characteristic of *A. baumannii* genomes. The genome plasticity of *A. baumannii* is notable (6) as it is able to take up DNA readily from its surroundings, thus increasing the likelihood of MDR.

Clinical epidemiology of *A. baumannii* is being investigated globally using multi-locus sequence typing (MLST), outer core locus (OCL) typing and capsule locus (KL) typing methodologies. MLST classifies isolates into clonal complexes based on allelic profiles of seven genes resulting in sequence type (ST) assignment (7), whereas OCL (8) and KL (9) typing use alleles of genes encoding the outer core assembly machinery and capsule production respectively. Tracking of clonal spread also aids in tracking ARGs and VGs associated with particularly concerning clones. However, more and more often, *A. baumannii* from non-clinical sources (10–13) are being interrogated because they too harbour ARGs and VGs. In fact, *A. baumannii* has been classified as a One Health problem (14), which encompasses all aspects of human, animal and environmental health. Therefore, it is critical to study clinical as well as non-clinical isolates of *A. baumannii* from many different sources in addition to global distributions.

Herein, a collection of 36 isolates from hospital, tank milk, agricultural surface water, stream, and waste-water effluent (WWE) highlights the incredible diversity of *A. baumannii* and substantiates that ARGs and VGs are not unique to hospital isolates and features similarities between clinical and non-clinical strains. Importantly, these isolates have been analyzed in the context of the main international clones described for *A. baumannii*, considering more than 149 additional genomes.

## Results

### Novel Sequence Types exist within isolates from non-clinical and clinical sources

There is great genetic and phenotypic diversity within *A. baumannii,* and our collection of isolates showcases this. Many of these isolates were characterized as novel STs (Pasteur MLST typing scheme) (7). The majority of novel STs (n = 13, 59%) were recovered from tank milk samples, (Figure 1a) from two countries: Germany and Indonesia. Notably, there were two novel STs from hospital sources: AB220-IK38 and AB224-IK42. The other novel STs were isolated from a stream (n=1), post-chlorinated-WWE (n=2), and agricultural runoff (n=4) (Figure 1a).

**Figure 1a:**
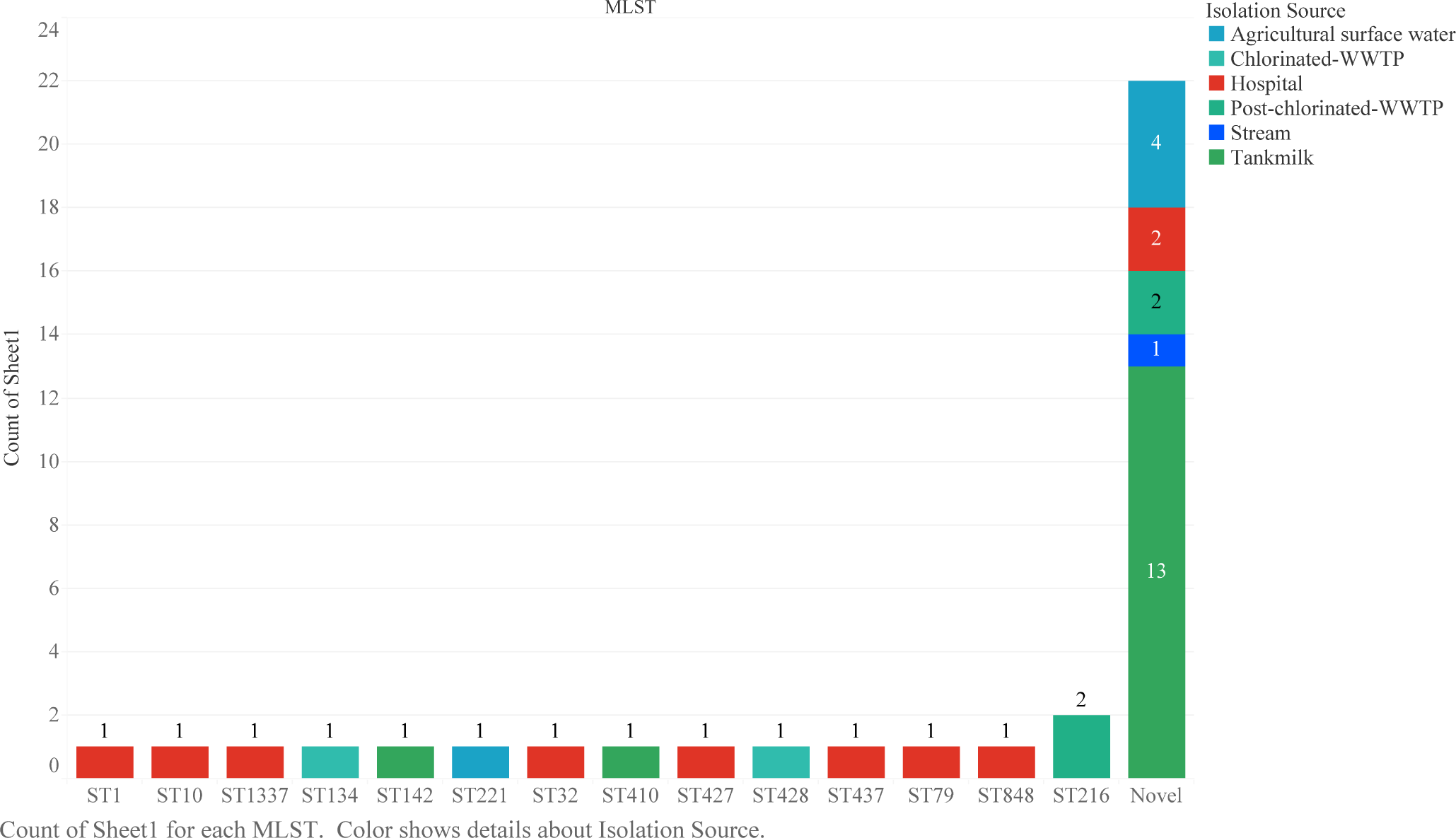
Diversity of sequence types in clinical and non-clinical isolate collection. Each strain in the collection was typed based on the Pasteur method of sequence typing. The number of isolates that fit into each ST category are shown. Those that are of a novel ST are not necessarily the same ST, but they are grouped into a novel category together to show the count. The colours indicate the isolation source of each isolate.

To further analyze this diversity, we examined the whole-genome sequences (WGS) using a phylogenomic approach and an additional 149 *A. baumannii* assemblies (Supplemental Table 2). AB337-IK11 and AB338-IK12, novel STs isolated from tank milk (Hesse, Germany), formed a new clade with closest relation to the most prevalent IC2 and with ATCC17978 (Figure 1b). The long length of the branch signifies this is a well-differentiated clade. Many of the human isolates with characterized STs clustered with IC1, IC5, IC6 and IC8. For example, one hospital isolate classified as IC1 (AB214-IK32; ST1) and another as IC8 (AB218-IK36; ST10).

**Figure 1b:**
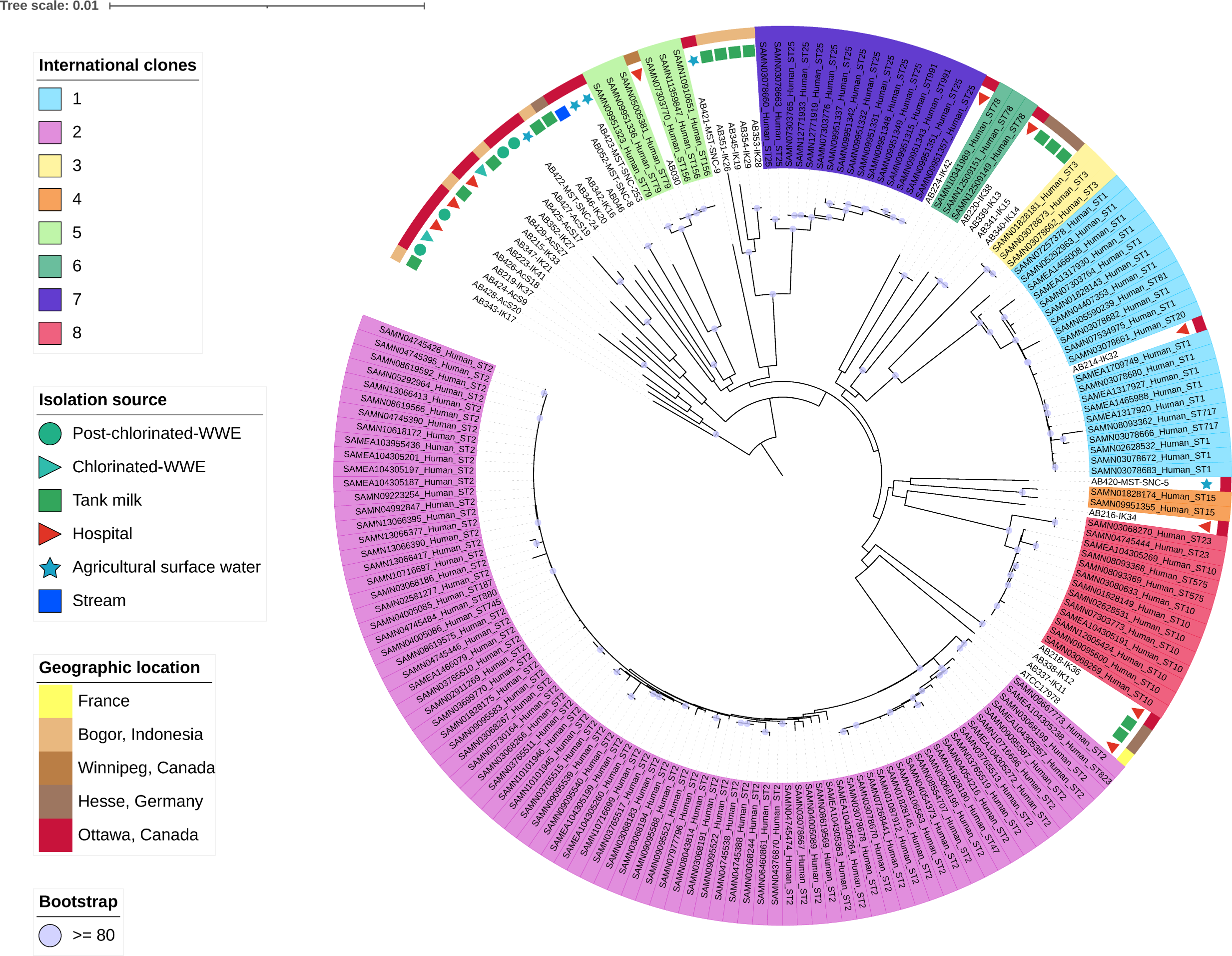
Phylogenomic analysis of isolates. Representative whole genome sequences from all eight international clonal (IC) groups were analyzed alongside the whole genome sequences of this collection. Condensed isolate names were used to ensure high readability. IC are signified by the large rectangles in the appropriate colour surrounding the assembly name. Isolation source is shown by the small shapes on the second most outer circle. The geographic origin is shown on the outer circle. Bootstrap values are indicated by a purple circle if they are > 80.

A highly diverse clade exists with closest relation to IC5 (Figure 1b) and is composed of 12 isolates. These isolates were recovered from Canadian hospital and post-chlorinated WWE samples and Indonesian tank milk samples. The following STs make up this clade; ST216, ST221, ST427, ST428, ST848 and ST1337 as well as novel STs. Interestingly, AB215-IK33, AB219-IK37 and AB223-IK41 from clinical samples, are distributed in this highly diverse clade. The detection of novel STs in the clinical and tank milk samples exemplifies the diversity of *A. baumannii* as does the lack of IC2 detection in our collection of clinical isolates.

### Non-clinical isolates show reduced susceptibility to clinically relevant antibiotics and their genomes contain antibiotic-resistance genes

Among this diverse set of isolates, antibiotic susceptibility profiles vary regarding clinically relevant antibiotics. AB030 and AB214-IK32, both from hospitals, showed MDR phenotypes based on CLSI breakpoints (Figure 2a). AB030 (15, 16) was used as a reference strain, in addition to ATCC17978 in this study. All isolates are susceptible to colistin. AB426-AcS18 from post-chlorinated WWE shows decreased susceptibility to cephalosporins (ceftriaxone, ceftazidime and cefepime) as well as the β-lactam antibiotic, piperacillin and its common β-lactamase inhibitor, tazobactam. The hospital isolate AB218-IK36 is resistant to doxycycline and ciprofloxacin. AB216-IK34 (hospital) displays resistance to ceftriaxone and ceftazidime. Another tank milk isolate, AB340-IK14 is resistant to ceftriaxone and a clinical isolate, AB215-IK33, displays resistance to meropenem with intermediate susceptibility to imipenem, piperacillin/tazobactam and ceftriaxone. Many of the non-clinical isolates (n=17, 46%) showed intermediate resistance to ceftazidime, ceftriaxone, piperacillin/tazobactam; all members of the β-lactam antibiotic family (Figure 2a).

**Figure 2a:**
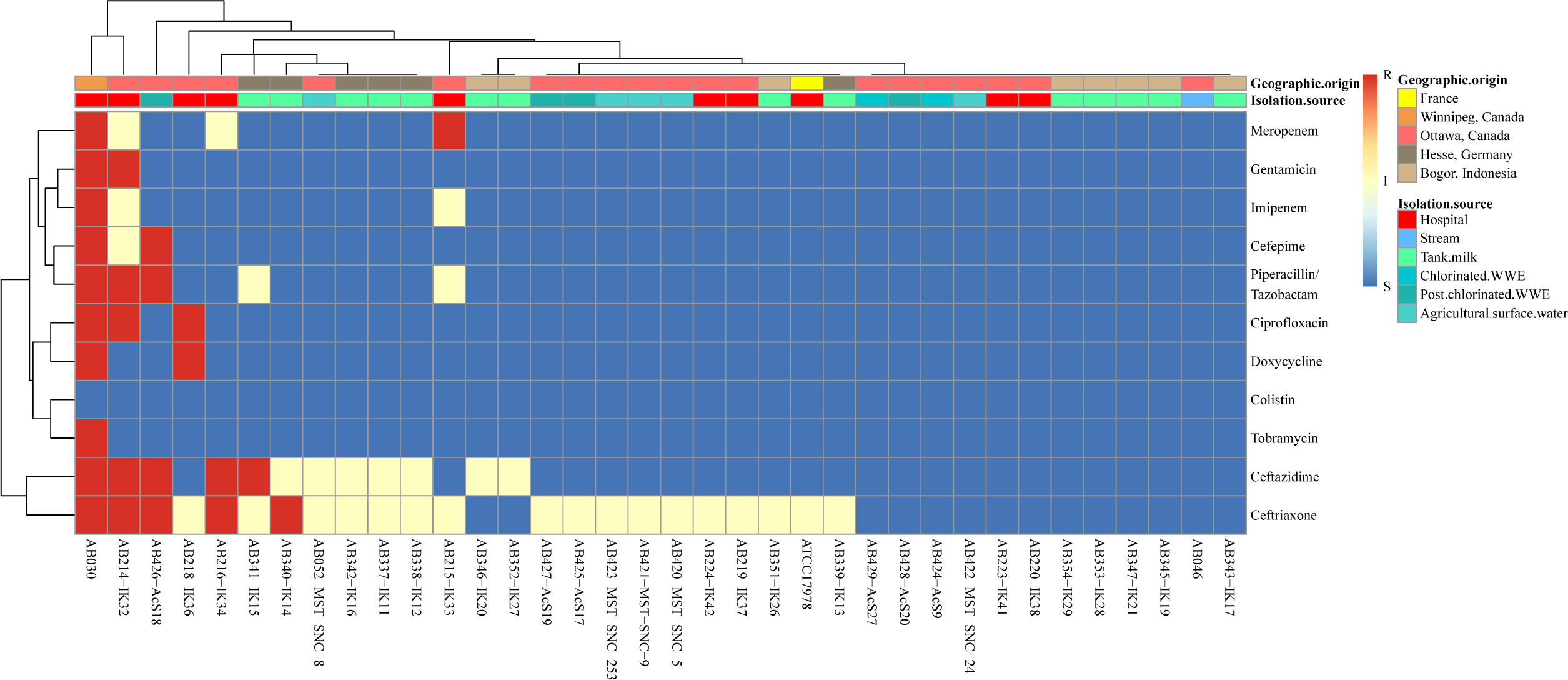
Susceptibility profiles of all strains. Susceptibility testing was performed according to the Clinical Laboratories Standards Institute (CLSI) guidelines. Clinically relevant antibiotics tested are part of the CANWARD panel and are as follows: amikacin, cefazolin, cefepime, cefoxitin, ceftazidime, ceftobiprole, ceftolazane/tazobactam, ceftriaxone, ciprofloxacin, clarithromycin, clindamycin, colistin, daptomycin, doxycycline, ertapenem, gentamicin, imipenem, linezolid, meropenem, nitrofurantoin, piperacillin/tazobactam, tobramycin, trimethoprim/sulphamethoxazole, and vancomycin. Only those antibiotics that showed at least one strain with a change in susceptibility based on CLSI breakpoints are shown.

The genetic contribution to susceptibility was then investigated. AB030 showed the most numerous genetic resistance mechanisms. Cephalosporinase encoding genes, (*bla* _ADC-5_, *bla* _ADC-16_, *bla* _ADC-18_) were scattered throughout the isolates, with *bla* _ADC-2_ being the most common (n = 8, 22%; Figure 2b). One of the non-clinical strains, AB429-AcS27, was found to harbour *bla* _ADC-79_. Despite the presence of this gene, AB429-AcS27 is susceptible to all clinically relevant antibiotics tested (Figure 2a). Finally, meropenem resistant AB215-IK33 (hospital, Ottawa, Canada), only shows the presence of *bla* _OXA-217_ and *bla* _ADC-5_. Aminoglycoside modifying enzyme encoding genes were rare; *aac(3)-Ia*, *ant(3’’)-IIa*, *aph(3’)-VIa* and *aph(3’)-Ia* detected in MDR strain AB214-IK32, and *aac(3)-IIe*, *aph(3’’)-Ib*, *aac(6)’-Ian* and *aph(6)-Id* in MDR strain AB030 (Figure 2b).

**Figure 2b:**
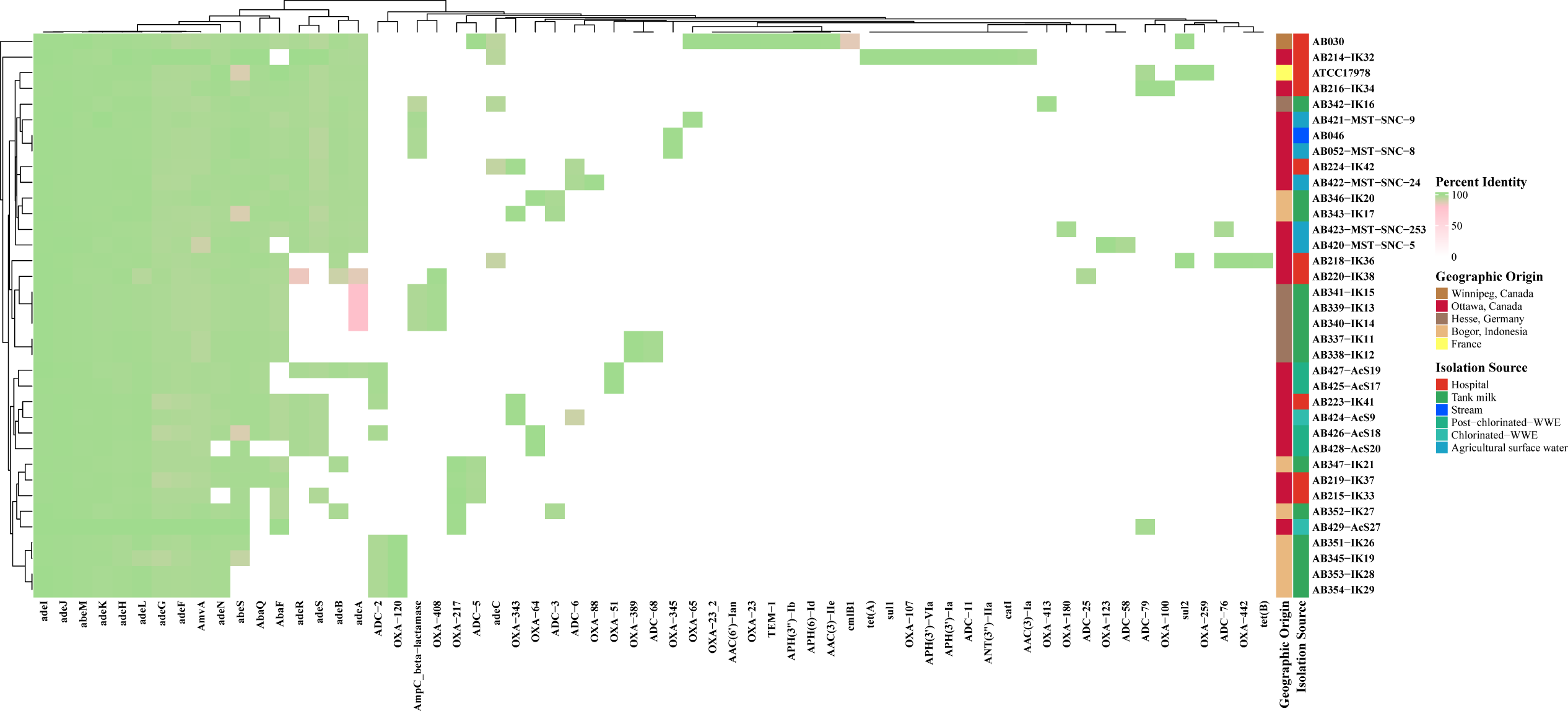
Antibiotic resistance gene profile of all isolates. Percent identity of each gene is used to explore the diversity. The minimum cutoff for nucleotide identity is 80% and is shown in pink. Hierarchical clustering demonstrate similarity based on the antibiotic resistant gene profile.

Efflux pump genes were highly conserved in all isolates. The genes encoding AmvA, a member of the MFS (Major-Facilitator Superfamily), and AbeM, an SMR (Small Multidrug Resistant) efflux pump, were strictly conserved and found in all isolates. Conservation of the RND (resistance-nodulation-division) family of efflux pumps is not a unique feature of *A. baumannii* (17) and these isolates are consistent with previous studies. Efflux operons *adeFGH* and *adeIJK* are found in all isolates, regardless of their geographic origin or isolation source. *adeAB* is the least conserved RND pump, with 49% (n = 18) of isolates encoding either *adeA* or *adeB.* Corresponding outer membrane factor, *adeC*, however, is much less conserved and is found only in 14% (n = 5) of isolates.

### Variation in efflux pump expression cannot only be explained by variation in promoter regions or regulators

Investigation into the expression of RND pumps in this collection of isolates was performed using the second gene in each operon as a proxy for expression. Within the clinical strains, AB030 displayed significant upregulation of all three characterized RND efflux pumps, while *adeIJK* was upregulated in two susceptible clinical isolates; AB219-IK37 and AB223-IK41 (Figure 3a).

**Figure 3:**
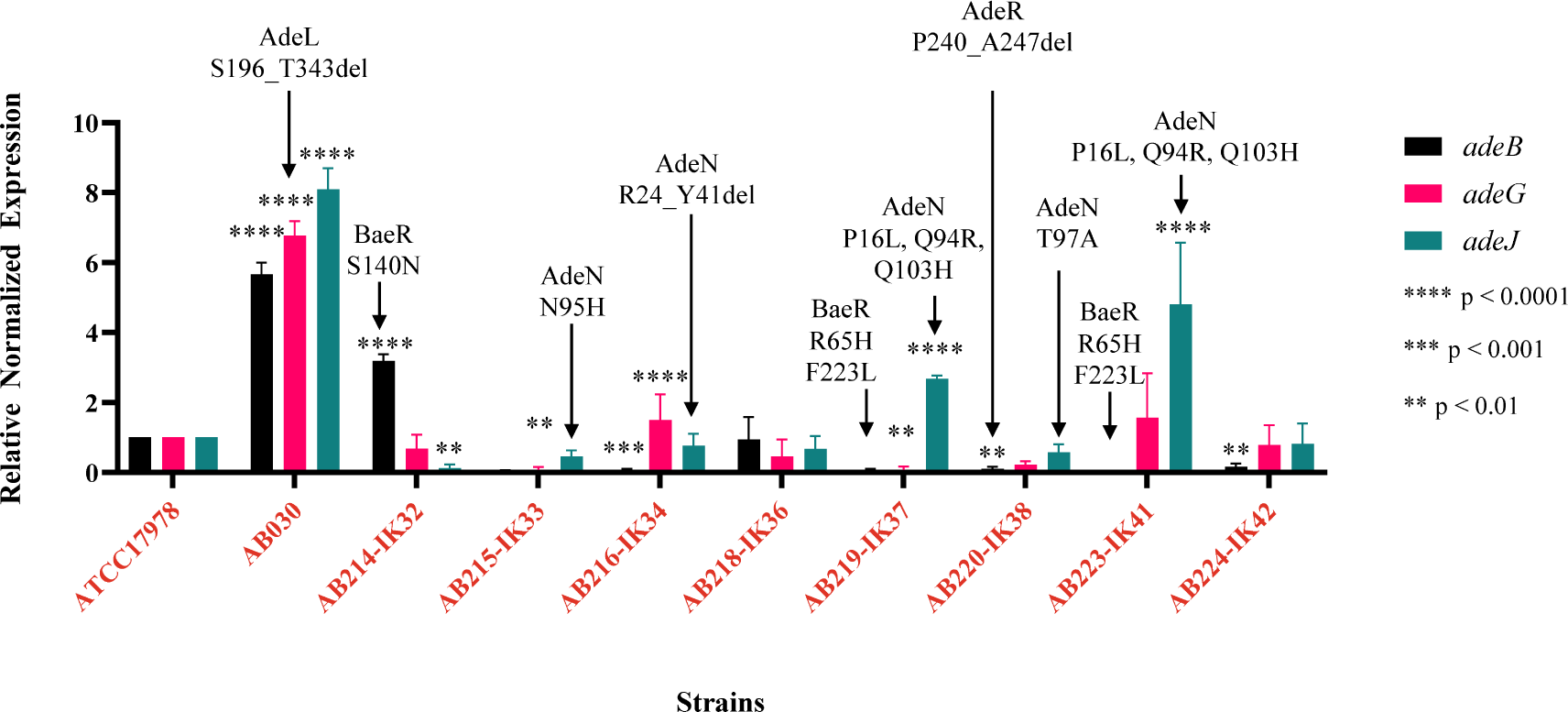
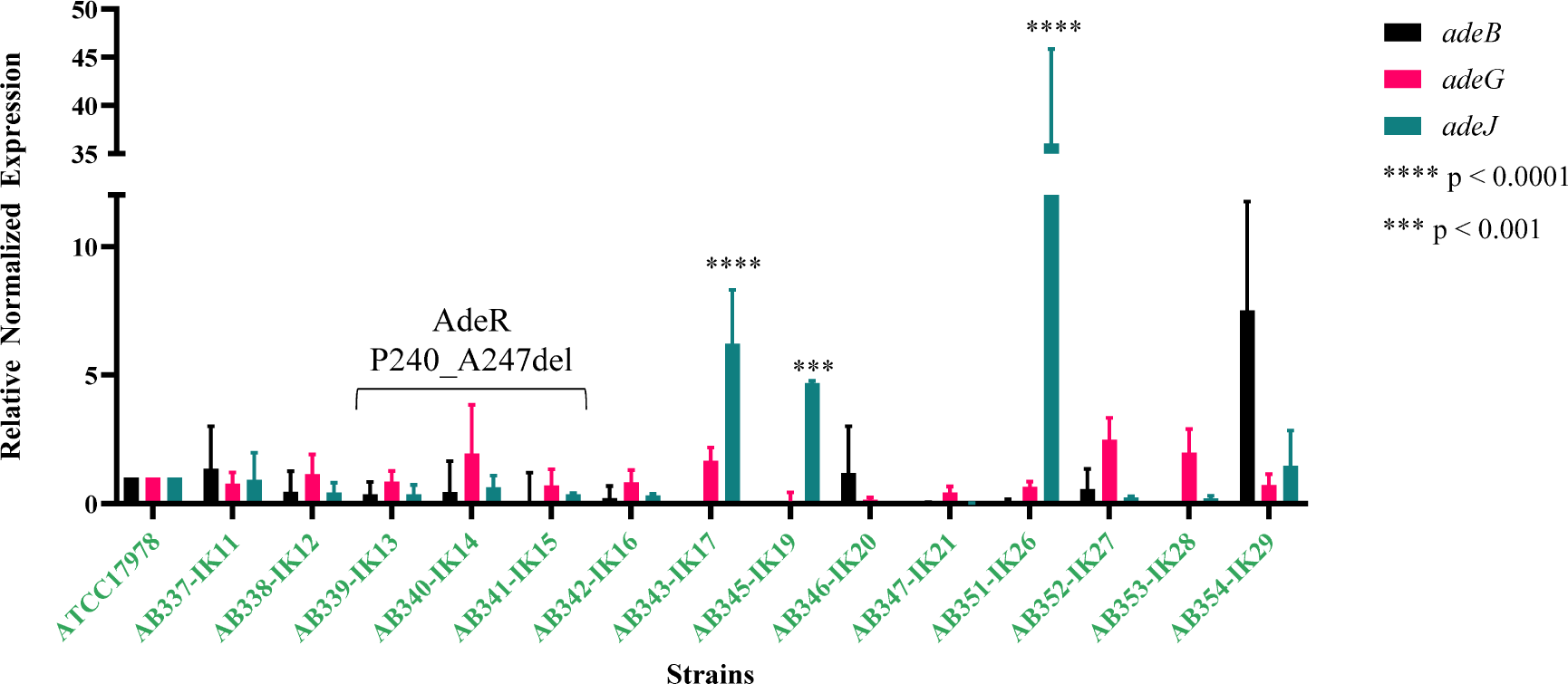
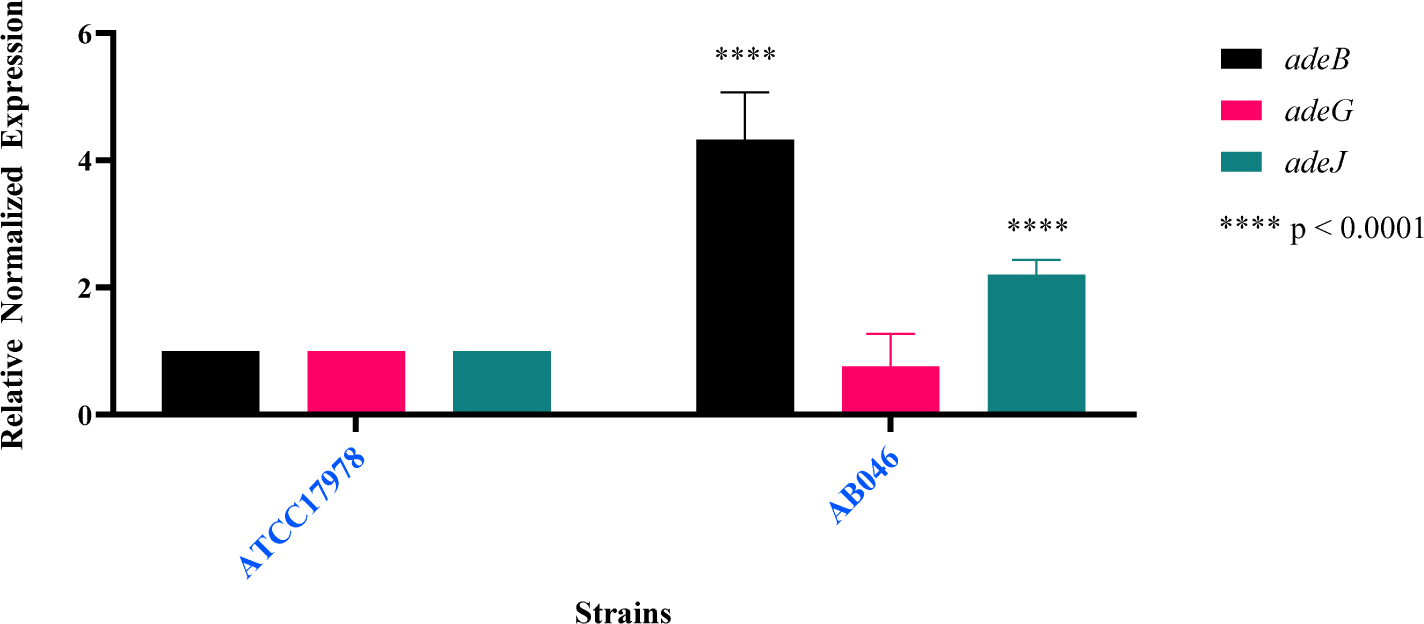
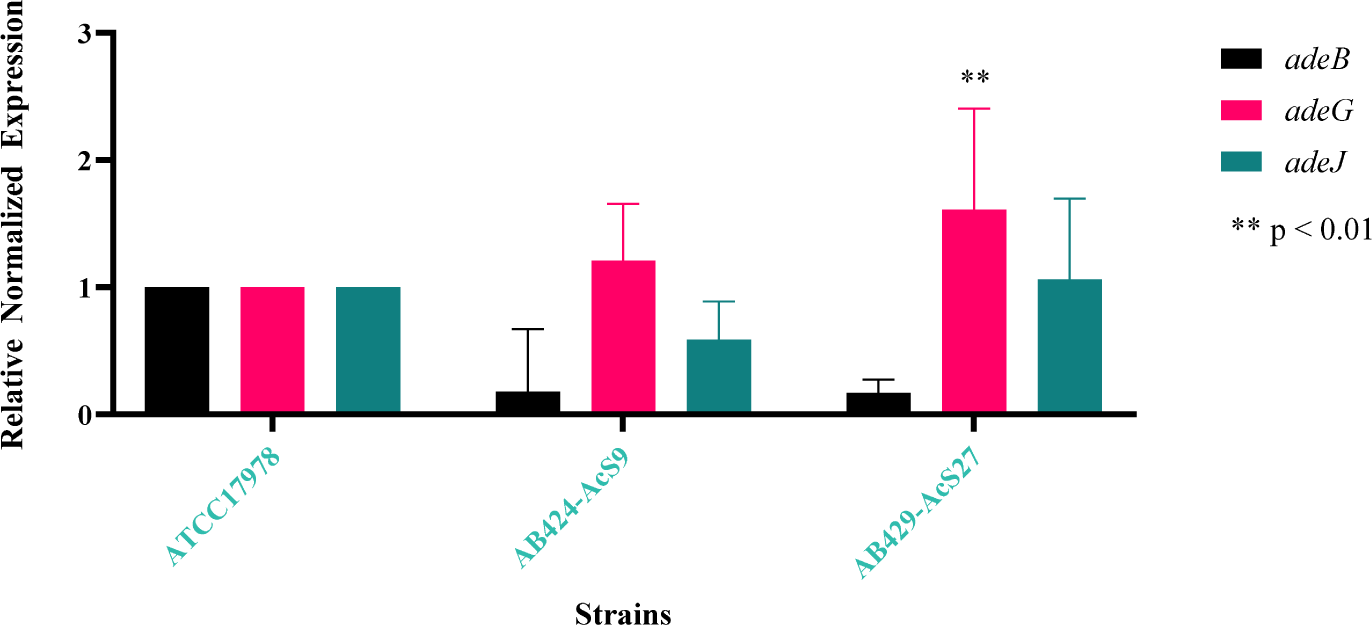
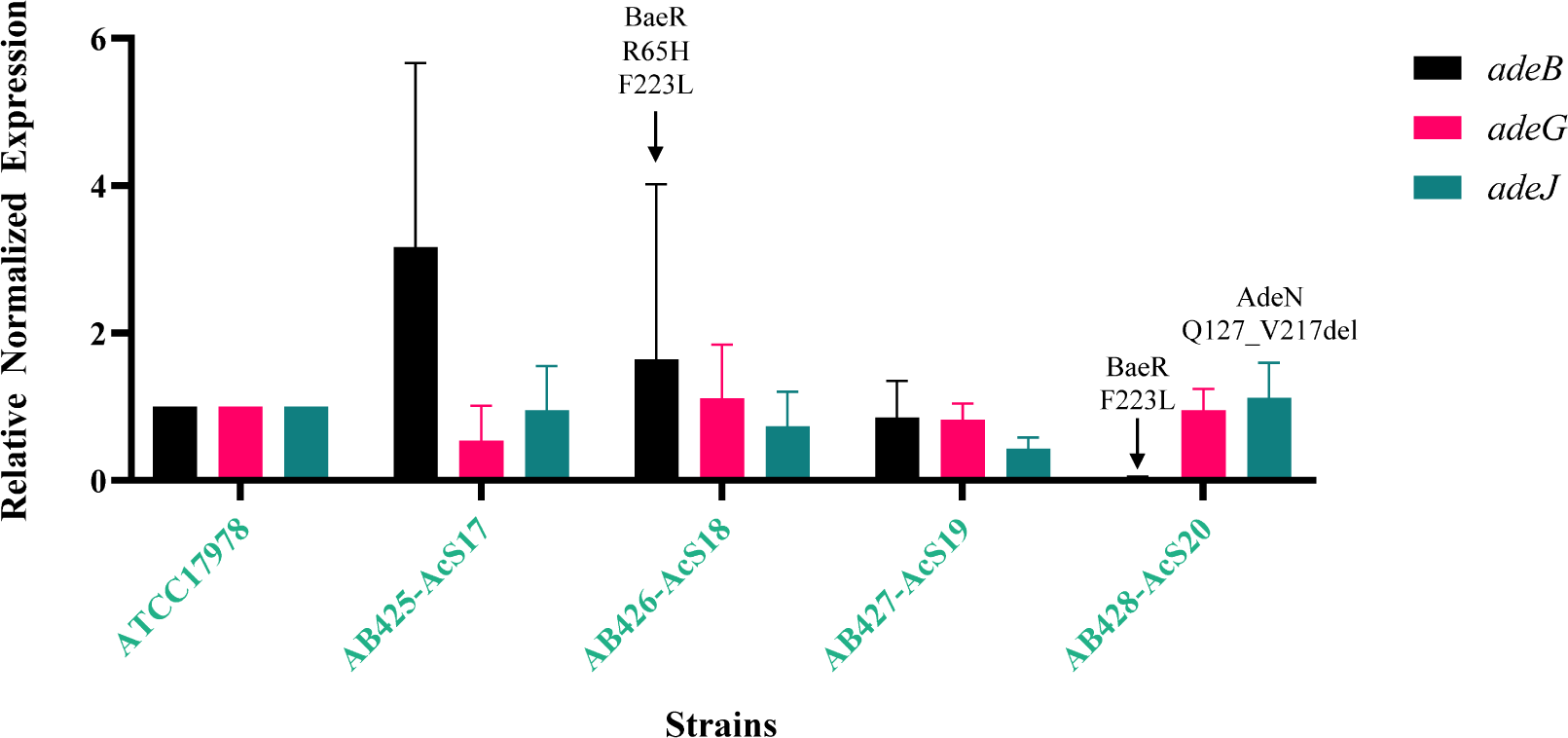
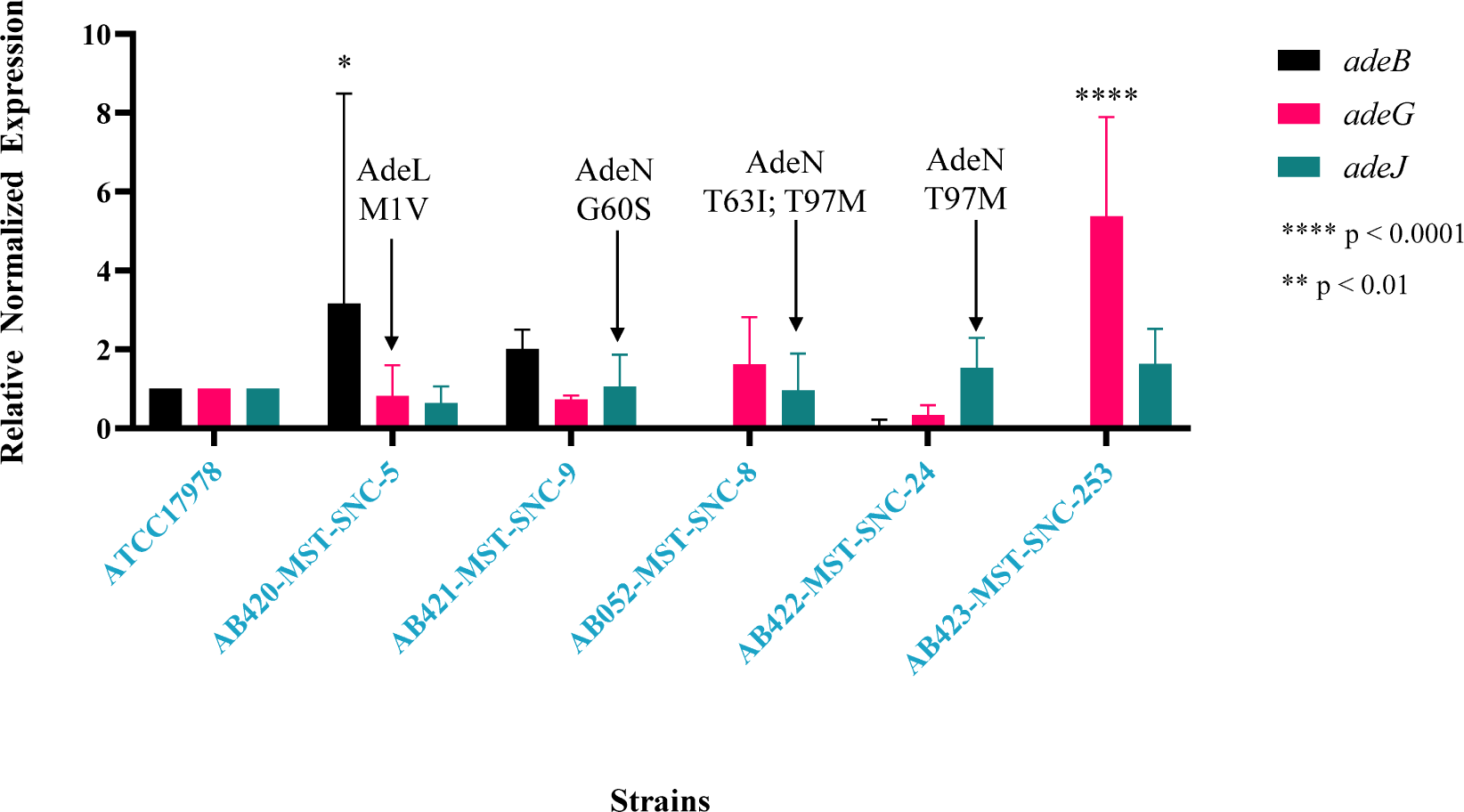
Resistance-Nodulation-division efflux pump expression of all strains. Expression was performed using total RNA converted to cDNA (see methods for more details) using primers listed in Supplementary table 1. Pfaffl method was used to calculate the relative fold change in expression compared to the type-strain ATCC17978. Each inset shows the clinical isolates, the stream isolate, the chlorinated and post-chlorinated-WWE isolates, and the agricultural surface water isolates. Two-way ANOVA was used in GraphPad 10.0.0 to determine significance. Significance value are indicated in each panel.

Overexpression of *adeIJK* was also observed in two isolates from tank milk, AB343-IK17 and AB351-IK26 as well as a stream isolate, AB046 (Figure 3b-c). The promoter region of *adeIJK* was investigated to determine possible explanations for this overexpression. It was 100% conserved between all isolates (data not shown) including the operator for AdeN, the TetR type regulator which exerts transcriptional control over *adeIJK* (18). Additional analysis was done for AdeN as well. Each isolate had 100% identity to the query sequence of AdeN, WP_000543879.1, except AB052-MST-SNC-8, AB428-AcS20, AB422-MST-SNC-24, AB220-IK38, AB215-IK33, AB421-MST-SNC-9, AB216-IK34, AB219-IK37 and AB223-IK41(n=9; 25%). None of these mutations were in the helix-turn helix DNA binding domain of these AdeN variants (data not shown) but most are found in the dimerization domain which is critical for AdeN function (Supplemental Figure 1a). AdeN of AB052-MST-SNC-8 contains T63I and T97M. AB428-AcS20 appears to have a truncation of AdeN at Q127_V217del (noted in Figure 3).AB422-MST-SNC-24 shares the same T97M as AB052-MST-SNC-8 while AdeN of AB220-IK38 shows T97A. There is a N95H mutation in AB215-IK33 and G60S mutation in AB421-MST-SNC-9 (Supplemental Figure 1a). In strain AB216-IK34, there is a deletion of 17 amino acids (R24_Y41del) and both AB219-IK37 and AB223-IK41 (hospital, Ottawa, Canada) have P16L, Q94R and Q103H mutations (Supplemental Figure 1b).

AB214-IK32 showed overexpression of *adeABC* (Figure 3a) which may contribute to its MDR phenotype (Figure 2a). The stream isolate AB046 also showed *adeAB* overexpression as did AB420-MST-SNC-5 (Canadian agricultural surface water) (Figure 3c, f). Hospital isolates AB216-IK34 and AB220-IK38 had a downregulation of *adeAB.* Of those isolates showing presence of *adeAB(C)* there was 100% nucleotide identity (data not shown). To investigate if the expression difference of AB214-IK32, AB220-IK38, and AB046 was due to possible regulator mutations, the two component systems, AdeRS and BaeSR were investigated.

No mutations appeared in AdeR of AB214-IK32, AB046, and AB420-MST-SNC-5. Further analysis of all isolates showed 100% conservation in the receiver domain, known to interact with its cognate sensor histidine kinase, AdeS (data not shown). However, the DNA binding domain of AdeR in AB220-IK38, AB339-IK13, AB340-IK14 and AB341-IK15 was truncated by 7 amino acids (Supplemental Figure 2a) while showing 100% identity to each other (Supplemental Figure 2b). Further to this, investigation into BaeR revealed a common mutation (R65H) observed in AB219-IK37, AB223-IK41 (both hospital, Ottawa, Canada) and AB426-AcS18 (post-chlorinated-WWE, Ottawa, Canada). AB214-IK32 from a Canadian hospital shows a S140N. Finally, a last group of strains displayed the F223L mutation including AB219-IK37, AB223-IK41 (hospital, Ottawa, Canada), AB426-AcS18, AB428-AcS20 (post-chlorinated-WWE, Ottawa, Canada), and AB353-IK28 and AB354-IK29 (tank milk, Bogor, Indonesia). The impact of these mutations is unknown and requires further study.

Regarding *adeFGH* expression, AB030 (hospital, Winnipeg, Canada) showed overexpression in agreement with previous studies (16), as did AB423-MST-SNC-253 (agricultural surface water, Ottawa, Canada) and AB429-AcS27 (chlorinated WWE, Ottawa, Canada). The promoter region of *adeFGH* which includes a portion of the reverse complement of its known regulator, *adeL*, as well as the AdeL regulatory site, showed 100% conservation for all isolates (data not shown). In AB030, however, there was a deletion of AdeL from S196 to the terminus at T343 (Supplemental Figure 3). As AdeL is a known repressor of *adeFGH*, this may provide an explanation as to why *adeG* is overexpressed in this isolate. Additionally, only one other isolate demonstrated variation in AdeL: The AB420-MST-SNC-5 mutation has an unknown impact on AdeL function (Supplemental Figure 3).

### Virulence is not associated with isolation source or geographic origin

Virulence is an important characteristic of pathogens and *A. baumannii* is no exception. The wax moth larvae insect model, *Galleria mellonella*, is a valid model to evaluate virulence in *A. baumannii* (19). Relative to ATCC17978, clinical isolates, AB215-IK33, AB216-IK34, AB219-IK37, AB220-IK38, and AB224-IK42 (n= 5, 41 % of clinical isolates) showed higher end point percent survival indicating less virulence (Figure 4a). AB218-IK36 was the only isolate showing higher virulence than ATCC17978. The remaining clinical isolates showed no significant difference and therefore were equally as virulent as ATCC17978. German tank milk isolate, AB342-IK16 and all Indonesian tank milk isolates showed higher survival rates (Figure 4b). AB046 from a Canadian stream, Canadian chlorinated-WWE isolates (n = 2) and three isolates from Canadian post-chlorinated WWE (Figure 4c) as well as AB052-MST-SNC-8, AB422-MST-SNC-24 and AB423-MST-SNC-253 from Canadian agricultural surface water (Figure 4d) showed less virulence.

**Figure 4:**
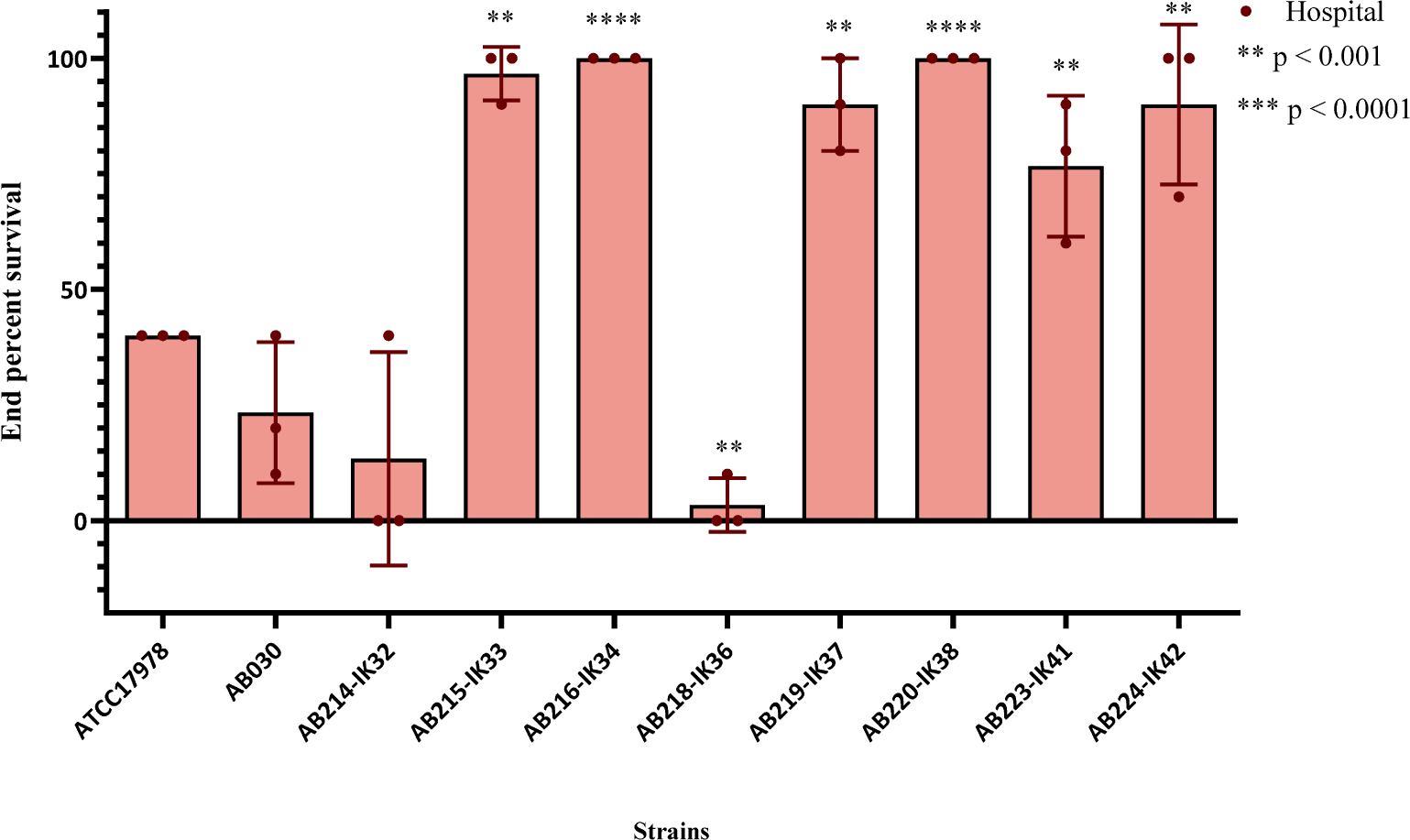
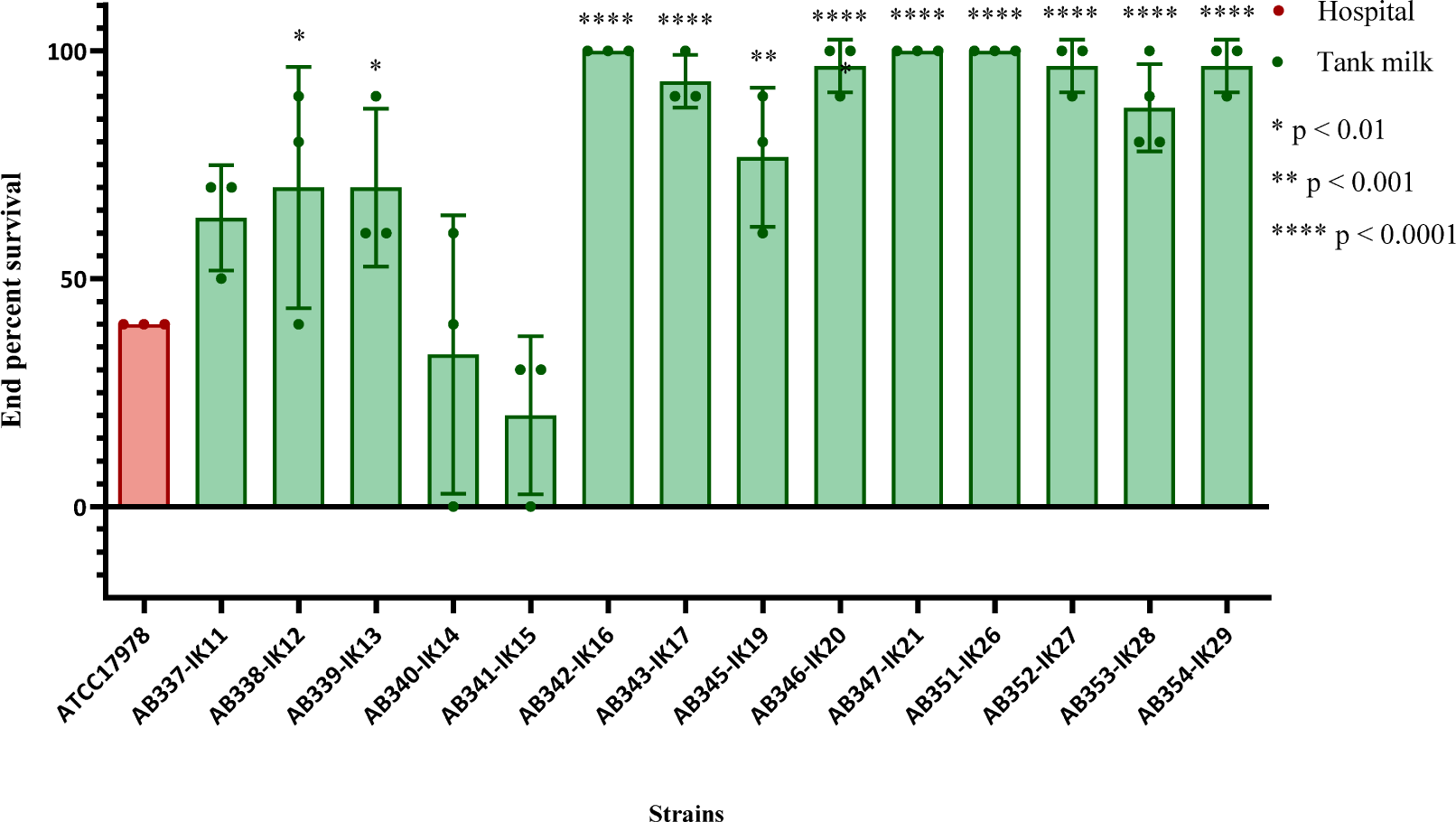
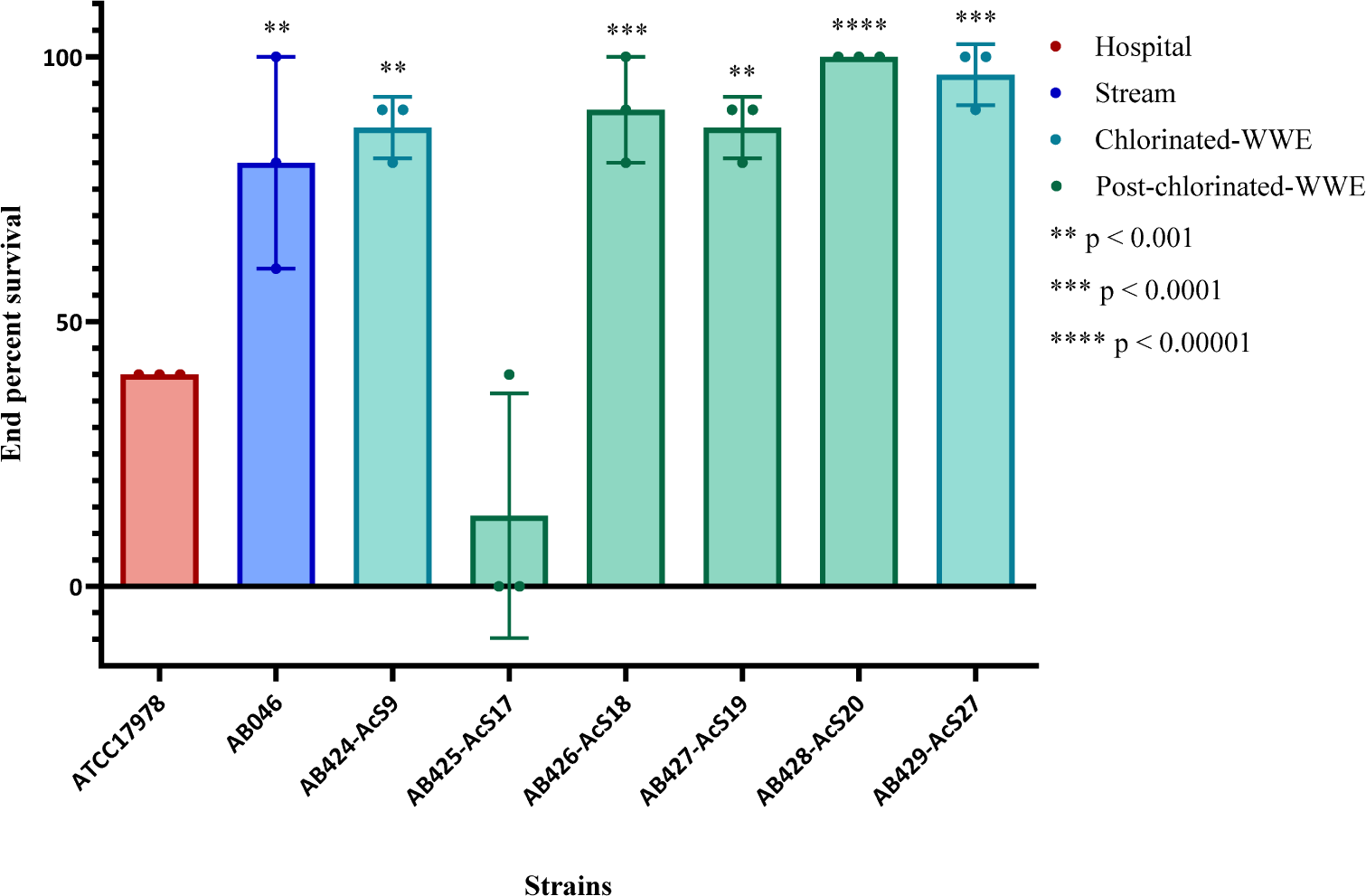
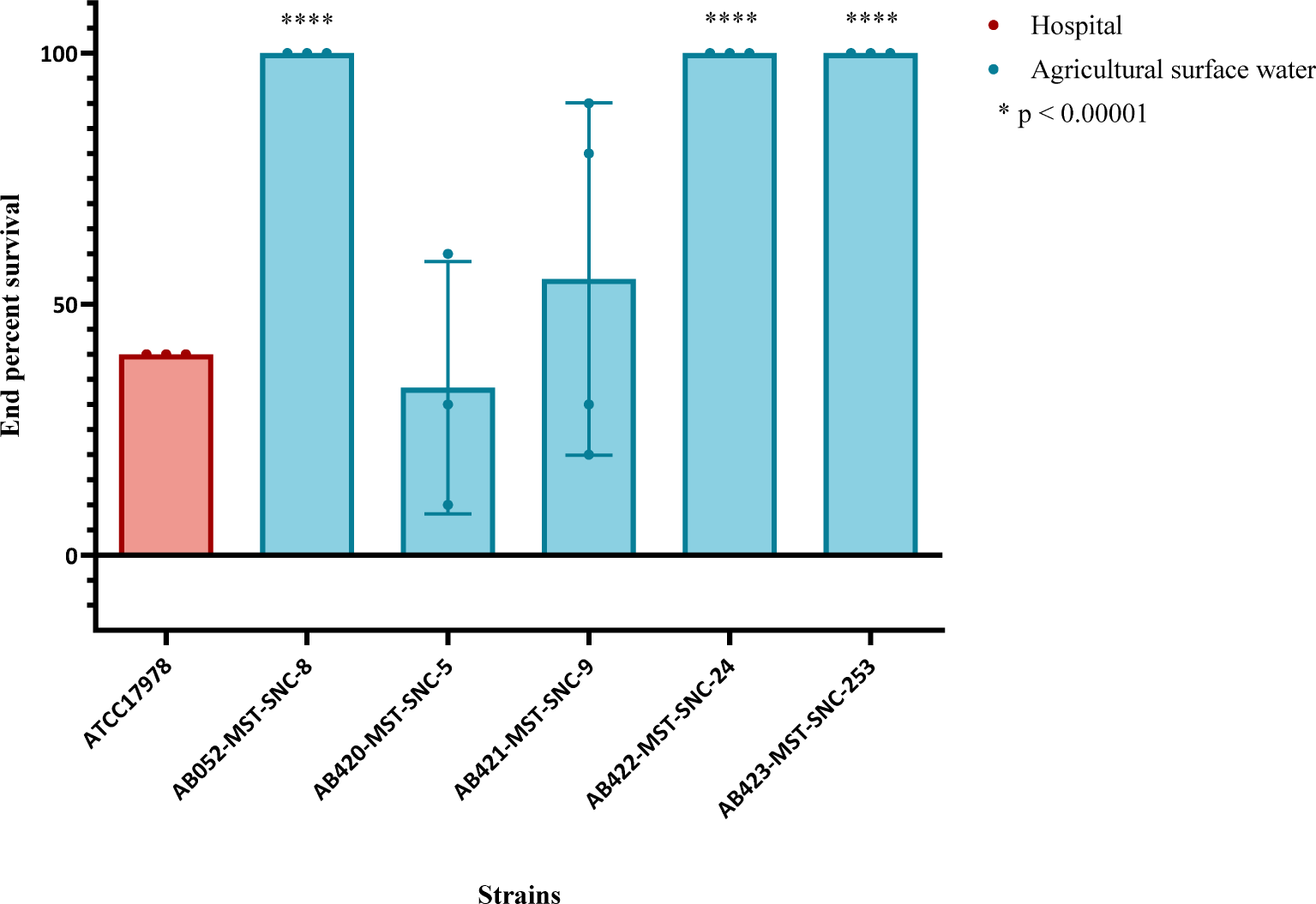
Virulence of hospital isolates in *Galleria mellonella*. Larvae were selected, wiped with 70% ethanol using a sterile swab and then injected in the back left proleg. Survival was measured every 12 hours for a total of 72 hours. Each biological replicate is represented as one point for each strain and is composed of 10 biological replicates for a total of at least 30 larvae for each strain. Statistics were performed in GraphPad 10.0.0. One-way ANOVA was performed to determine significance. One star indicates p<0.1. Two stars indicates p< 0.01, three stars indicate p<0.001. Four stars represents p≤0.0001. **A**: Virulence of hospital isolates from Ottawa, Canada. **B**: Virulence of tank milk isolates, from Bogor, Indonesia and Hesse, Germany. ATCC17978 is included for comparison despite being a hospital isolate and is therefore indicated in red. **C**: Virulence of the stream isolate, the chlorinated-WWE and post-chlorinated-WWE isolates, all from Ottawa, Canada. ATCC17978 is included for comparison despite being a hospital isolate and is therefore indicated in red. **D**: Virulence of agricultural surface water isolates. ATCC17978 is included for comparison despite being a hospital isolate and is therefore indicated in red.

Hospital isolates from Ottawa, Canada clustered together with tank milk isolates from Bogor, Indonesia and post-chlorinated WWE isolates from Ottawa, Canada (blue ellipsis, Figure 5) based on principle component analysis (PCA). These isolates have similar virulence profiles suggesting virulence is independent of geographic origin and isolation source. Further, in a separate cluster, German tank milk strains grouped with clinical isolates from Ottawa, Canada (green ellipsis, Figure 5). There are also those strains whose virulence profiles are distinct from any other strain in the collection, for example, the type-strain ATCC17978 (red circle, top right, Figure 5). Clusters of clinical and non-clinical strains highlight overlap in virulence between these strains from widely different geographic origins and isolation sources.

**Figure 5:**
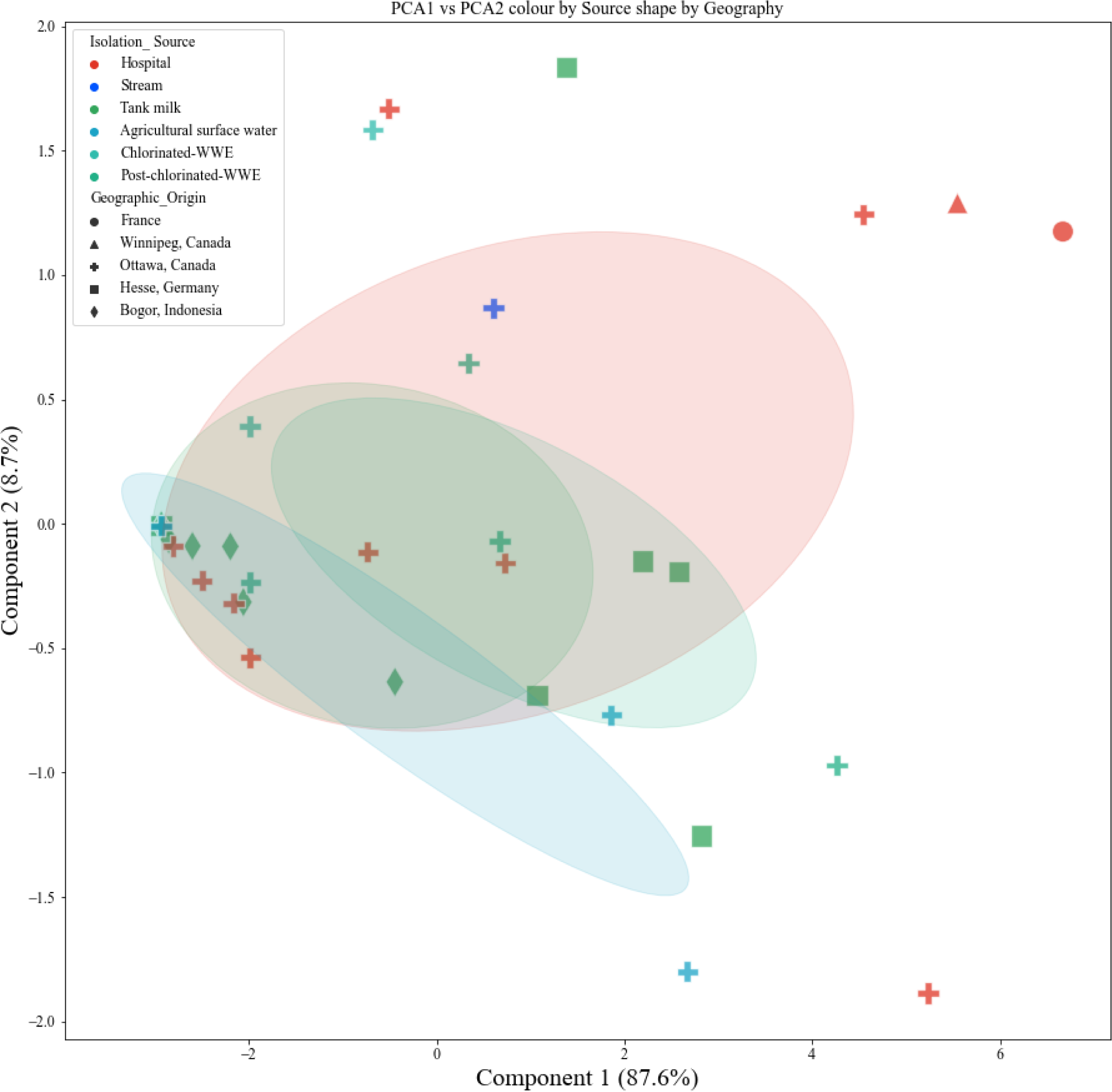
Principal component analysis of survival of *G. mellonella*. Survival data was used to determine if clinical and non-clinical isolates clustered together. Isolation source is indicated in various colours and geographic origin is noted as varying shapes. Ellipses include area that is 1 standard deviation from the mean with 95% confidence intervals. Ellipses colours match the isolation source colour that corresponds to the cluster.

### The presence or absence of virulence genes does not explain virulence profiles in *G. mellonella*

In order to understand the genetic components behind the virulence profiles observed, the VGs of all strains were investigated. Non-clinical and clinical isolates alike, regardless of geographic origin, cluster together as there are similarities in VG profiles (Figure 6).

**Figure 6:**
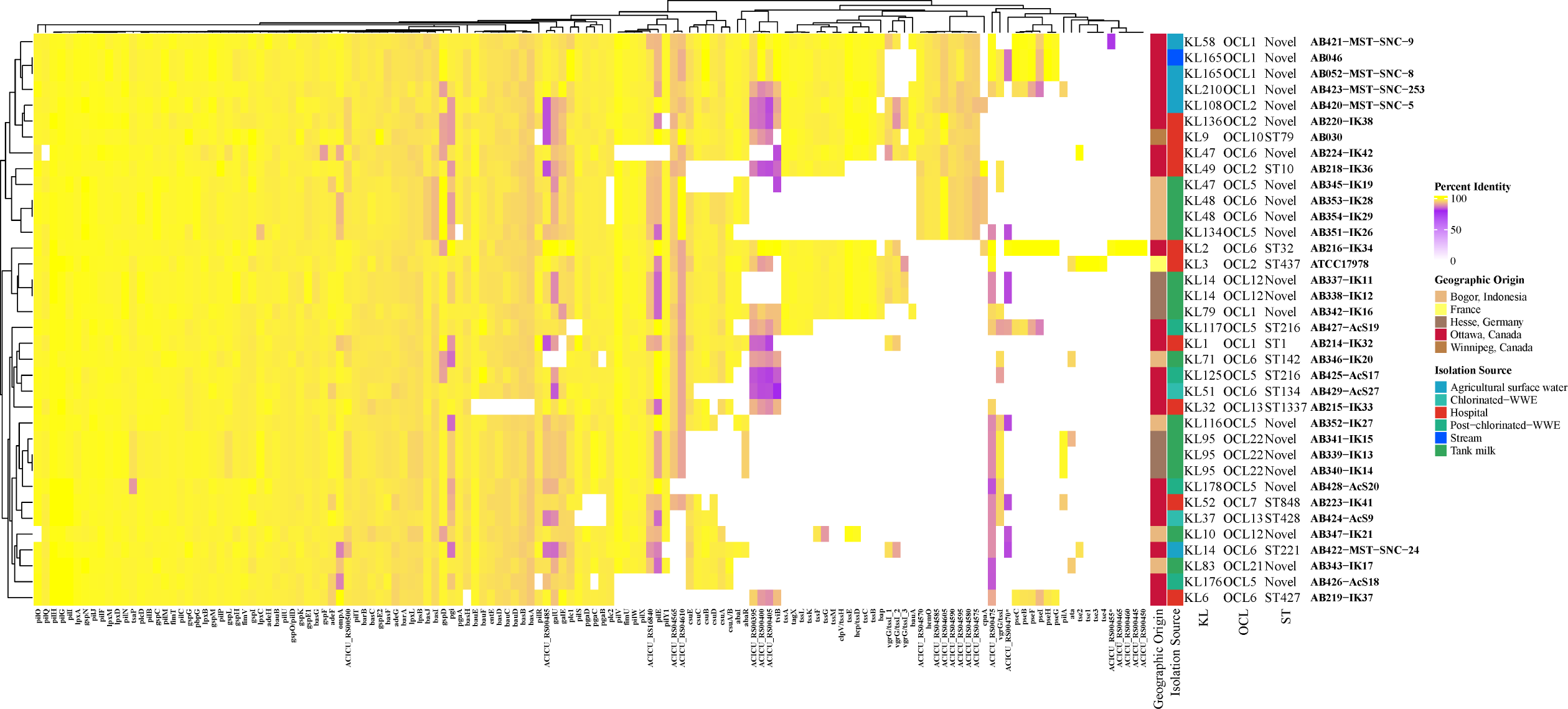
Virulome determination of all isolates. Percent identity of each gene is used to compare the diversity. The minimum cutoff for nucleotide identity is 80% and is shown in purple. Hierarchical clustering demonstrate similarity based on the virulence gene profile. Capsule locus (KL), outer core locus (OCL) and Sequence Type (ST) are listed for each isolate.

The outer-membrane encoding gene, *ompA*, was conserved in all isolates (Figure 6). The glycoprotease gene, *cpaA*, was detected in hospital (AB216-IK34, and AB218-IK36), in tank milk (AB346-IK20, AB354-IK29), and agricultural surface water (AB420-MST-SNC-5, AB421-MST-SNC-9 and AB422-MST-SNC-24) isolates. There was also high conservation of the *pil* genes involved in production, secretion and assembly of the Type 4 pili (T4P) used in surface attachment and twitching motility. There was most variation in *pilE* which encodes a minor pilin subunit and has not been well characterized in *A. baumannii*. Further, only AB224-IK42, AB339-IK13, AB340-IK14, AB341-IK15, AB342-IK16 and AB423-MST-SNC-253 have alleles of the major pilin subunit, *pilA* above 80% identity. Interestingly, ATCC17978 had the highest motility and appeared to be associated with forming lower amounts of biofilm (Supplemental Figures 4ad and 5a-d). This was interrogated further in all isolates via (Spearman’s) correlation analysis that was not significant (R = −0.1382, p = 0.4016); thus, no correlation between biofilm and motility was observed (Supplemental Figure 6).

Typing and epidemiological tracking of *A. baumannii* and other Gram-negative pathogens not only consists of MLST but also of OCL and KL typing. OCL and KL typing also provide a starting point for determining lipopolysaccharide (LOS) and CPS structures. The majority of these isolates were typed as OCL6 (n = 8, 22%) including AB224-IK42 (hospital). In our collection, associated STs with OCL6 are ST32, ST142, ST134, ST221, ST427 and 3 novel STs. Six (17%) isolates are OCL1, including isolates from clinical and non-clinical sources. The OCL2 designation was assigned to three isolates, including the type-strain ATCC17978, AB420-MST-SNC-5, from chlorinated-WWE and AB220-IK38, from hospital. A wide distribution of KL types was observed. This includes the most common KL14 and KL95 (each n = 3, 8%). Those KL types associated with at least one isolate of a novel ST include: KL14, KL10, KL47, KL48, KL58, KL79, KL83, KL95, KL108, KL116, KL134, KL136, KL165, KL176, KL178, and KL210 (Figure 6). The most common clinical KL type is KL9 (20) and only AB030 (hospital, Canada) was identified as KL9, further supporting the unexplored diversity of *A. baumannii*.

## Discussion

Overall, the extensive diversity of *A. baumannii*, both genetic and phenotypic is highlighted in this study. This collection encompasses a distinct set of human and non-human isolates from diverse isolation sources and geographic origins. Detection of novel STs from clinical and non-clinical sources exemplifies this non-uniformity. Despite IC2 being the dominant IC in Canada (21), none of these hospital isolates were categorized as such. As shown in Figure 1b, one hospital isolate, AB214-IK32 was classified as an IC1, which is the second most prevalent Canadian clonal complex. AB218-IK36 (ST10) is IC8 which was also previously detected in Canada (21). The lack of relation of the clinical isolates to those already characterized in Canada suggests clonal replacement of other lineages, at least in Ottawa, Canada, since the most recent study in 2019 (21).

AB337-IK11 and AB338-IK12 (tank milk, Germany) are unique because they cluster between IC1 and IC8 and form their own well-differentiated clade (Figure 1b). WGS phylogenetic characterization indicates that they belong to the same ST, albeit novel. This conclusion is further supported by their KL and OCL types; both strains are KL14 and OCL12 (Figure 6). This collection contains isolates of the most recently identified OCL types – OCL17 - 22(22); one isolate from Indonesian tank milk; AB343-IK17 (OCL21) and three from OCL22 namely AB341-IK15, AB339-IK13 and AB340-IK14, all from German tank milk and all with novel STs (Figure 6). ST32 has previously been associated with OCL6 (22), whereas our study shows that ST142, ST134, ST221, ST427 and 3 novel STs have now been associated with this OCL type. AB337-IK11 and AB338-IK12 also reveal intriguing hierarchical clustering based on VG analysis. Not only do they cluster together but they also cluster with clinical isolate ATCC17978 (Figure 6) as they did in the phylogenomic analysis (Figure 1b) and both do not show a significantly different virulence profile compared to ATCC17978 (Figure 4b). These two isolates highlight the diversity of non-human strains. The distribution of hospital isolates throughout the diverse clade near IC5 also exemplifies these dissimilarities between WGS of clinical and non-clinical strains (Figure 1b).

This collection of *A. baumannii* shows great variation in susceptibility, ARG profile, virulence in *G. mellonella* and VG profile. Hierarchical clustering suggests that there is no clear-cut trend with regard to susceptibility and isolation source or geographic origin (Figure 2a). The genes *bla* _ADC-79_ *, bla* _ADC*-*3_ and *bla* _OXA-217_ were detected in AB429-AcS27 (Canadian chlorinated-WWE). Tight control of β-lactamase expression is maintained and only induced in the presence of the β-lactam drug (23) suggesting that the presence of the antibiotic alone should induce expression and therefore confer resistance. However, AB429-AcS27 is susceptible to all antibiotics tested, including all β-lactam antibiotics, signifying alternate expression control in this strain. Another strain that showed presence of *bla* _OXA-217_ was AB215-IK33 (Canadian hospital). In contrast to AB429-AcS27, this isolate is resistant to meropenem and upon further investigation, both *bla* _ADC-5_ and *bla* _OXA-217_ genes were detected. The *bla* _OXA-217_ is a *bla* _OXA-51-like_ β-lactamase gene and although not specifically characterised, members of this group are known to hydrolyze imipenem and meropenem (24). Therefore, it is likely responsible for this phenotype in AB215-IK33. Insertional elements, such as IS*Aba*1 are known to increase expression of *bla* genes such as *bla* _OXA-23-like_, *bla* _OXA-51-like_, *bla* _OXA-58-like_, and *bla* _OXA-235-like_ genes (24). Further work determining expression control of *bla* _OXA-217_ in AB215-IK33 (resistant to meropenem) and AB429-AcS27 (sensitive to meropenem and other antibiotics tested) will be critical to understanding these phenotypes in non-clinical isolates. These two strains, among others, exemplify the crucial need to test susceptibility in addition to investigation of the ARGs. Additionally, the detection of *bla* in non-clinical *A. baumannii* is not unique to this study (11, 25, 26). However, the detection of these elements in *A. baumannii* from non-clinical sources such as water and soil suggests that they act as reservoirs for ARGs, especially β-lactamases (24) and this analysis supports this.

Overexpression of RND efflux pumps is a major contributor to MDR in clinical *A. baumannii* (27), however, these pumps are not always expressed. Five strains showed overexpression of *adeIJK* without presenting a MDR phenotype (Figure 3). Identified substrates of AdeIJK include β-lactams, chloramphenicol, tetracycline, erythromycin, lincosamides, fluoroquinolones, fusidic acid, novobiocin, rifampin, and trimethoprim (28). AdeIJK is known as the universal *Acinetobacter* spp. efflux pump and is typically constitutively expressed (29), therefore, this overexpression is surprising. AB219-IK37 (hospital, Canada), and AB351-IK26 (tank milk, Indonesia) are intermediately resistant to ceftriaxone while AB223-IK41 (hospital, Canada), AB343-IK17 (tank milk, Indonesia) and AB046 (stream, Canada) are susceptible to all antibiotics tested including those substrates of AdeIJK tested. Analysis of AdeN of AB052-MST-SNC-8, AB422-MST-SNC-24, AB421-MST-SNC-9 (all agricultural surface water, Canada), AB428-AcS20 (post-chlorinated WWE, Canada), AB215-IK33, AB219-IK37, AB220-IK38 and AB223-IK41 (all hospital, Canada) revealed a common variance at T97 (n=2: T97M and n=1: T97A), as well as at P16L, Q94R and Q103H (n=2), N95H, G60S, T63I and a truncation at Q127_V217del (Figure 3) all located in the dimerization domain (Supplemental Figure 1). Dimerization of TetR-type repressors, including AdeN, is essential for function. None of these mutations are in the the helix-turn-helix DNA binding domain of AdeN, however. The P16L, Q94R, Q103H mutations in both AB219-IK37 and AB223-IK41 could explain the overexpression of *adeIJK* in these strains. This does not however explain the susceptibility of these strains to those substrates of AdeIJK (Figure 2a). Considering there is no overexpression of *adeIJK* in any of the other strains with AdeN mutations (Figure 3), t this needs further investigation with additional sequencing and characterization. The truncation of AdeR in AB220-IK38 (hospital, Ottawa, Canada) may be attributing to the decreased expression of *adeB* but further experimentation is required.

As antibiotic selective pressure is higher in clinical settings, we predicted that those isolates from clinical isolation sources would display higher levels of resistance. This was not the case, however. Two hospital isolates, AB220-IK38 and AB223-IK41 were susceptible to all antibiotics tested and clustered with those of non-clinical origin (Figure 2a). Vice versa, AB426-AcS18 (post-chlorinated WWE) clustered with hospital isolates. In almost all cases, hierarchical clustering created different groupings based on ARG and susceptibility profiles. It is notable that AB030 and AB214-IK32 clustered together (Figure 2a) as having the most similar susceptibility profiles, as well as the most similar ARG profile (Figure 2b). This is also despite being from different ICs (AB030; IC5 and AB214-IK32; IC1) and different geographic origins (AB030; Winnipeg, Canada and AB214-IK32; Ottawa, Canada). A susceptibility-based cluster includes AB052-MST-SNC-8 (Canadian agricultural surface water), AB342-IK16 (German tank milk), AB337-IK11 (German tank milk), and AB338-IK12 (German tank milk) (Figure 2a) whereas AB052-MST-SNC-8 clusters with AB224-IK42 (Canadian hospital) in the ARG analysis (Figure 2b). A hospital isolate and an agricultural surface water isolate clustering together due to similar ARG profiles but clustering separately based on susceptibility phenotypes is an intriguing finding. One short-coming of this study is that expression for every ARG was not determined, and this may explain some of the differential clustering between ARG and susceptibility profiles. However, these data provide additional support for the need for phenotypic testing of resistance instead of genetic analysis alone.

There is a bias in the literature towards virulence characterization using an animal model for only those isolates from hospitals (30, 31). To our knowledge, only two non-clinical isolates have been investigated for virulence (10, 32). Therefore, we investigated if the virulence profiles of our hospital isolates were comparable to those from non-clinical sources. The majority of the clinical isolates were less virulent compared to ATCC17978 (Figure 4). This collection of isolates represents strains from non-dominant ICs, such as IC1, IC4, IC7 and IC8, which broadens our understanding of the diversity of *A. baumannii* virulence. Interestingly, the clinical isolates were undifferentiated from those of non-clinical sources (Figure 5). The lack of delineation between clinical and non-clinical isolates suggests that isolation source is not a good indicator of virulence. It also suggests that a non-human isolate of *A. baumannii* could just as easily become a pathogen if given the opportune conditions.

It is intriguing that a high variation in *pilA* is observed. PilA is the major subunit of the T4P and there is evidence of convergent evolution with other bacterial species (33). Modification of pilins via glycosylation supports host immune evasion (34) and in a similar fashion, variation or lack of canonical *pilA* may also provide immune avoidance strategies. Furthermore, in other studies, there appears to be an inverse correlation between biofilm formation and motility (35) in a PilA-dependent manner (33). However, our data does not corroborate this finding (Supplemental Figure 6). When performing a Spearman correlation analysis between motility and biofilm biomass, we did not find a correlation (R = - 0.1382, p = 0.4016). This may be due to the fact that our study included non-human *A. baumannii* where previous work was done in clinical isolates.

It is interesting that *cpaA* was detected in non-clinical isolates. CpaA has been characterized in clinical isolates of *Acinetobacter* spp. as a key player in hydrolysis of human *O*-linked glycoproteins whose cleavage results in blood coagulation (36, 37) and complement disruption (38). To our knowledge, the characterization of such a potent virulence factor has been limited to clinical isolates, and therefore this presents an interesting avenue for further study. As a potent glycoprotease, only known to cleave human *O-*linked glycoproteins, the conservation of *cpaA* in those isolates from the non-clinical origins suggests alternate functions or conservation of surrounding genetic elements that are advantageous. Furthermore, this also highlights the pathogenic potential of *A. baumannii* from non-clinical sources.

Many non-human bacterial isolates harbour ARGs as a means to protect themselves from other antibiotic producers in their environmental communities (39). To truly understand the origin of antibiotic resistance and virulence traits in *A. baumannii,* isolates from all environments must be characterized, as well as the microbial communities surrounding them including antibiotic producers and their genes (40). Studies have shown that ARGs identified in soil-dwelling microbes have been mobilized to clinical isolates (41).

In conclusion, these results show that by exploring the diversity of *A. baumannii* genetically and phenotypically, the prevalence of ARGs in non-clinical isolates becomes even more apparent and that these non-clinical isolates act as reservoirs for their clinical counterparts. Very few *bona fide* VGs are attributed to *A. baumannii,* where mechanisms to resist desiccation, withstand harsh conditions and evade host immune defences are hallmarks of successfully pathogenic strains (34). We also demonstrate high variability of VGs requiring investigation for the development of vaccines and classification of novel VGs and alleles. Our study greatly contributes to the variation and wide distribution of ARGs and VGs in *A. baumannii* especially in non-human isolates. Finally, in order to combat the ever-evolving infections caused by *A. baumannii*, a One Health approach must be taken. Non-clinical strains harbour ARGs and VGs, may be an important genetic reservoir for clinical isolates and represent the unexplored diversity of *A. baumannii*.

## Materials and Methods

### Strain Isolation

A summary of the strains isolated for this study can be found in Supplemental Table 1.

### Whole-Genome Sequencing

Detailed methods for extraction of DNA and sequencing can be found in the Supplemental Methods. All assemblies have been deposited to NCBI GenBank with the following: Bioproject PRJNA819071, and Biosamples SAMN26898552 - SAMN26898587.

### Species Determination, Phylogenomic Analysis, Sequence Typing, Outer Core Locus Typing and Capsule Typing

The average nucleotide identity (ANI) was calculated with pyani v.0.2.9 (42) using Mummer 3.0 (43) to confirm the species using the *A. baumannii* strain ATCC 19606 (Biosample: SAMN13045090). Only isolates above 95% of identity with the reference strain were considered for further analysis. The phylogenomic analysis was performed as previously described (44). In brief, a pangenome analysis was run by means of roary (45) using 95% identity as the cutoff value. The genes in a single copy in all the genomes, commonly known as single gene families (SGFs), were recovered from the pangenome matrix. Then, the SGFs were aligned and tested for recombination. A maximum likelihood phylogeny was built with 483 SGF without recombination, and finally, the tree was annotated. Assemblies used can be found in Supplemental Table 2. Sequence Typing was performed using the Pasteur scheme (7) as reference via ABRicate (46). Outer core locus and capsule typing were performed using Kaptive (9).

### MIC Determination

Antibiotic susceptibility was evaluated as per the CLSI guidelines using broth microdilution methods. The full CANWARD panel (47) of antibiotics was used for testing. Based on CLSI breakpoints, isolates were categorized into susceptible (S), intermediate (I) and resistant (R) (48). MIC values can be found in Supplemental Table 3. The data was displayed using pheatmap in R Studio.

### Antibiotic Resistance Gene Determination

The ABRicate pipeline (46) was used and directed to use the Comprehensive Antibiotic Resistance Database (CARD) (49) for analysis of all genomes with perfect (100% identity) and strict (≥80% identity) with hit coverage of 80% or greater. Output was then displayed using the ComplexHeatmap package (50) in R Studio.

### RND Efflux Expression Evaluation

Overnight cultures of each strain were subcultured 1/100 in LB broth with shaking at 37°C until mid exponential phase (A_600_ of 0.7±0.5). Cells were pelleted and supernatant removed before storing at −70°C overnight. RNA was extracted using the Invitrogen Purelink RNA extraction kit (Thermo Fisher, Waltham, USA), according to the manufacturer’s protocol. After which, 1 µg of eluted RNA was treated with 2 U of DNase using the Invitrogen Purelink DNase kit (ThermoFisher, Waltham, USA) and incubated at 37°C for 30 minutes followed by inactivation at 62°C for 5 minutes. cDNA was synthesized using the Invitrogen VILO cDNA kit (ThermoFisher, Waltham USA) from 1 µg DNase treated RNA. Incubation conditions were as follows: 25°C for 10 minutes, 42°C for 60 minutes, and then 85°C for 5 minutes. The newly synthesized cDNA was diluted 1/5 prior to analysis using RT-qPCR. Applied Biosciences (Beverly Hills, USA) SYBR select master mix was combined with the appropriate volume of sterile mQH_2_O and primers to a final concentration of 200 nM. Primers sequences can be found in Supplemental Table 3. Diluted cDNA template was added so that 200 ng of DNA is in each reaction. Reaction was run on an Applied Biosciences (Beverly Hills, USA) StepOnePlus system with the following thermal cycling parameters: 50°C for 2 minutes, 95°C for 2 minutes, then 40 cycles of 95°C for 15 seconds, 60°C for 1 minute, with a final melt curve stage of 95°C for 10 seconds, 60°C for 5 seconds, and a step up to 95°C for 10 seconds increasing temperature by 0.3°C. RT-qPCR for each strain was performed with three technical replicates and at least three biological replicates per target gene. The Pfaffl method of analysis was used to calculate the relative fold change expression compared to ATCC17978. One-way ANOVA was performed using GraphPad Prism 10.0.0.

### Virulence Gene Determination

The ABRicate pipeline (46) was directed to use the Virulence Finder Database (VFDB) (51) to analyze all genomes with ≥ 80% nucleotide identity minimum and hit coverage of ≥ 80%. Output was then displayed using the ComplexHeatmap package (50) in R Studio.

### Survival Assays with *Galleria mellonella*

Upon arrival, larvae were sorted based on size and all those that were deemed equal were used. Larvae were injected within one week of arrival on site. Overnight cultures of *A. baumannii* were standardized to 0.5 MacFarland and then diluted 1/100 in 0.85% sterile saline. The ventral side of each larva was swabbed with 70% ethanol in preparation for injection. Then 10 µL of standardized diluted culture was injected into the left hind proleg. Larvae were then incubated in sterile plastic petri plates at 37°C for 72 hours. Survival was measured every 12 hours. At the beginning of each injection session, sterile 1x PBS was injected into 10 larvae as well as at the end to ensure syringe washes were adequate. Ten technical replicates and at least three biological replicates were performed for each strain. Percent end-point survival was calculated, and One-way ANOVA in GraphPad Prism 10.0.0 was used to evaluate significance compared to ATCC17978. PCA was performed in Python using Pandas and visualized using Seaborn and Matplotlib.

## Acknowledgements

The authors would like to thank Nancy Laing for assistance with susceptibility testing. A special thanks to Adam Sykes for his expertise in Python and principal component analysis. We also thank Dr. David R. Lapen and his field crew (AAFC), and Dr. Marc Desjardins (Ottawa Hospital Research Institute) for providing surface water samples and clinical isolates as well as co-op students for lab assistance.

The funders had no role in study design, data collection and interpretation, or the decision to submit the work for publication. This study was funded by a Discovery Grant from the Natural Sciences and Engineering Council of Canada to AK. IUHK is funded by grants from Agriculture and Agri-Food Canada under Biological Collections Data Mobilization Initiative (BioMob, Work Package 2; J-001564), A-base Fungal and Bacterial Biosystematics (J-002272) and Managing AAFC’s biological collections (J-002295) projects.

## Author contributions

Ellen M. E. Sykes - Data curation, Formal Analysis, Investigation, Methodology, Project Administration,, Visualization, Writing – original draft, Writing – review & editing

Valeria Mateo-Estrada - Data curation, Formal Analysis, Visualization, Writing – review & editing

Raelene Engelberg – Investigation, Writing – review & editing

Anna Muzeleva – Investigation, Writing – review & editing

George Zhanel - Methodology, Writing – review & editing

Jeremy Dettman - Investigation, Writing – review & editing

Julie Chapados - Investigation, Writing – review & editing

Suzanne Gerdis - Investigation, Writing – review & editing

Ömer Akineden – Investigation

Izhar UH Khan – Conceptualization, Funding Acquisition, Methodology, Writing – review & editing

Santiago Castillo-Ramírez - Conceptualization, Funding Acquisition, Methodology, Writing – review & editing

Ayush Kumar - Conceptualization, Funding Acquisition, Methodology, Resources, Supervision, Writing – review & editing

